# How many gamebirds are released in the UK each year?

**DOI:** 10.1101/2020.10.22.350603

**Authors:** Joah Robert Madden

## Abstract

Large numbers of gamebirds (pheasants *Phasianus colchicus*, red-legged partridges *Alectoris rufa* and mallard *Anus platyrhynchos*) are released annually in the UK to support recreational shooting. It is important to know how many of these birds are being released because their release and management has ecological effects on the wildlife and habitats of the UK. There is little regulation governing their release and consequently the numbers being released are unknown. I made 71 estimates of the numbers released based on numbers being reported formally via import controls and the Poultry Register, or extrapolated based on the breeding outputs of reared birds, or extrapolated based on the likely numbers and behaviour of shoots, or extrapolated based on observations of surviving birds. Based on the set of these estimates that fall within credible boundaries (ranging from 14.3 to 70.1 million birds), I estimate a mean of 34.5 million birds (95%CI 30.9-38.1 million) and a median value of 32.1 million (IQR 22.0-44.9 million) being released. This suggests that 24.3-25.3 million pheasants, 4.2-9.4 million partridges and 1.0-4.9 million mallard are released annually in the UK. These figures are markedly lower than previous published estimates and I discuss why such differences may occur. I set these figures in the context of the number and behaviour of shoots operating in the UK.

## INTRODUCTION

Recreational hunting (aka shooting) in lowland UK is predominantly based on the harvest of birds that have been specifically reared and released each year for the purpose (Martin 2011, 2012). Pheasant *Phasianus colchicus* and Red-legged partridges *Alectoris rufa* are not native to the UK but have been introduced as quarry species on several occasions over the past centuries and have become naturalised (Lever 1977). Mallard *Anas platyrhynchus* are native and their breeding and release for shooting is believed to have started in 1890 (Sellers & Greenwood 2019). It is estimated and commonly stated that tens of millions of individuals of these three species are released annually and an index of numbers being released has shown increases since the early 1960s of around nine times for pheasants (Robertson et al. 2017), just over five times for mallard (Aebischer 2013) and almost 200 times for red legged partridges (Aebischer 2013). However, the most recent estimate of the numbers of pheasant and red-legged partridge involved is accompanied by very large margins of error (being between 38-49% of the mean values). For pheasants, Aebischer (2019) calculated that 47 million birds were released annually, with 95% confidence intervals spanning 39 to 57 million. For red-legged partridges, Aebischer (2019) calculated that 10 million birds were released annually, with 95% confidence intervals spanning 8.1 to 13 million. There was no estimate of the numbers of mallard released. The only published mallard release numbers date back to 1985 when around 500,000 were estimated to be released (Harradine 1985)

An accurate understanding of the number of birds being released is important. The release of large numbers of birds and their subsequent management is believed to cause a variety of positive and negative ecological impacts on the habitats and wildlife of the UK (reviewed by Madden & Sage 2020, Sage et al. 2020). Each year’s release introduces a large amount of biomass into the ecosystem (Blackburn & Gaston 2018). Releases usually occur in lowland UK during the late summer, with cohorts of birds being placed in woodland release pens (pheasants), farmland pens (partridges) and ponds (mallards). The direct effects, caused by the birds themselves, are typically bad for habitats and wildlife in the immediate areas where releases happen and might be expected to scale with the numbers of birds released in a rather predictable manner (Sage et al. 2005). At high densities, the released birds may exert effects including physical disturbance of soils, altering nutrient levels, changes in floral composition and changes in invertebrate community composition (Reviewed in Madden & Sage 2020, Sage et al. 2020). The birds disperse from the release pens over the following months and during this time they are commonly killed or scavenged by generalist predators including foxes and raptors (Madden et al. 2018). The carcases provide a nutrient resource and it has been suggested that they support higher numbers of predators than might naturally be expected (Roos et al. 2018). These releases are accompanied by associated effects on habitats and wildlife that are the result of human management motivated by gamebird release, including retaining, planting or managing areas of cover vegetation, the control of predators deemed a threat to the released birds, and the provision of supplementary feed. Unlike direct effects, and the relationship between the numbers of birds released and the occurrence or scale of associated effects is not likely to be so straightforward (Reviewed in Madden & Sage 2020, Sage et al. 2020). In general, bird release is accompanied by larger areas of woodland being established or maintained, higher levels of predator control and large quantities of supplementary feed is provided either via hoppers or seed-bearing crops (Reviewed in Madden & Sage 2020, Sage et al. 2020). In order to better understand the scale of the direct and associated effects accompanying the release of birds for shooting, it is essential to have a reliable figure of the numbers of birds being released.

It would appear that the Poultry Register, administered by the Animal and Plant Health Authority (APHA) should be able to provide a definitive number of birds being released. Prompted by the growing recognition of risks to poultry and wild birds from avian influenza, a registration system was established in 2006 as a requirement of the Avian Influenza (Preventative Measures) (England) Regulations 2006. This Register is obligatory for holdings with flocks of more than 50 birds and voluntary registration of flocks that are smaller than 50 birds is encouraged. It covers the whole UK, explicitly includes birds kept for ‘restocking game birds’, and demands detailed numerical and spatial data about how many birds of what species are held where and for what purposes. It asks for separate numbers of birds of each species that are held for breeding for shooting, rearing for shooting and released for shooting. The APHA Poultry Register records for 2019 (obtained under an FOI request on 29 Jan 2020 with a response on 13 Feb 2020) show that: 10,039,379 pheasants and 3,819423 partridges and 435,907 duck (species unspecified) were reported as held for release for shooting. This indicates that a total of 14.3 million birds (of all species) are released annually. These figures are 21-38% of the totals estimated by Aebischer (2019) and fall well outside the confidence intervals that he presented. This disparity between the two current best estimates of release sizes suggests that at least one of the two current measures is likely to be very inaccurate.

Alternative estimates of the numbers of birds being released may be possible by conducting a series of extrapolations based on other current and historical datasets that relate to their rearing, release, management and shooting (Fig 1). First, indications of the numbers of birds that are reared annually may be available from import figures relating to the numbers of chicks or eggs brought into the country. It may also be possible to calculate the number of eggs or chicks being produced in the UK based on the anticipated productivity of adult birds reported as being held for breeding. Second, non-regulatory, economic surveys of shoots may provide data on the numbers of birds that are being released on shoots of different types and sizes. By extrapolating these numbers while accounting for the size and type of shoots, estimates may be obtained. Third, released birds require intensive management and so by considering the area over which the birds might be released and the numbers of gamekeepers available to manage them as well as the numbers of birds that a keeper may be able to manage, estimates may be obtained. Fourth, many shoots are commercial entities and so advertise the numbers of birds that they offer to be shot per day. Using these figures in conjunction with numbers of days when shooting occurs and incorporating measures of harvesting efficiency, estimates may be obtained. Fifth, individual hunters (known as and hereafter referred to as ‘guns’) may record the numbers of birds that they shoot and when considered in conjunction with numbers of guns, perhaps divided into classes relating to the scale and regularity of their shooting, estimates may be obtained. Finally, released birds that survive to the following breeding season may be counted as part of national Breeding Bird Survey (Woodward et al. 2020). By incorporating mortality rates (both natural and from harvest) for released birds, it is possible to back-calculate how many birds might have had to be released in order that the observed populations might be surveyed. All six approaches (like the published work of Aebischer (2019) and the poultry register data) are probably flawed with each one relying on partial data that may not be representative. Consequently, I have adopted a multilateration approach in which I search for consistencies or inconsistencies across the eight methods of estimates and crucially, ask how those estimates correspond to the reported behaviour of game shoots in terms of the numbers of birds that an ‘average’ shoot reports releasing and shooting.

**Figure 1.**
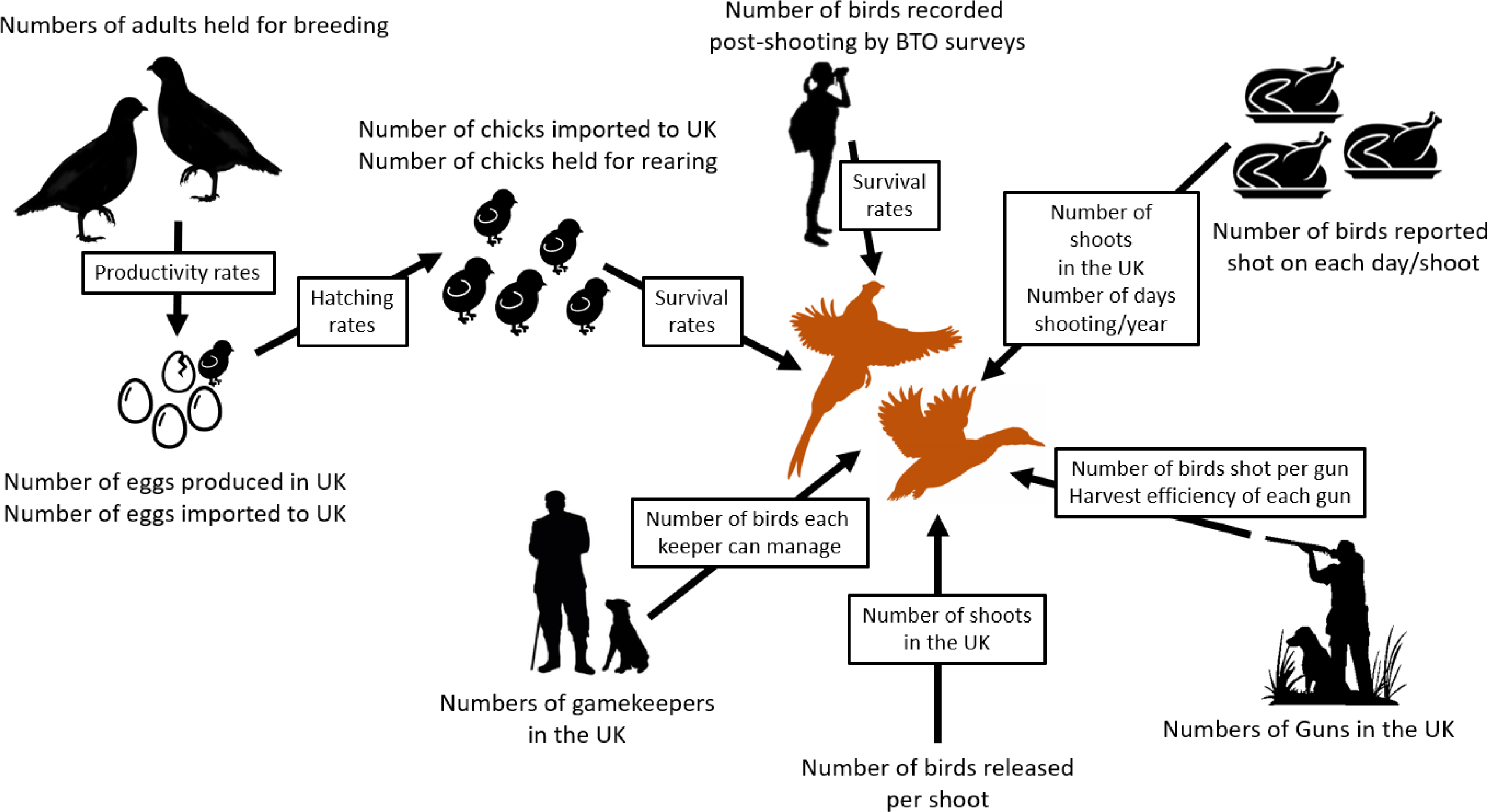
The various ways that datasets may be combined in order to generate estimates of the numbers of gamebirds released in the UK.

## METHODS

### Overview

I drew on a variety of data sets to supply measures of: numbers of birds reared; numbers of birds being released; numbers of days when shooting occurred; harvest sizes; number of people that shoot; and numbers of birds observed in the wild. I then multiplied various permutations of these measures in order to generate estimates of numbers of released birds (see Results and Table 1).

**Table 1.** The set of the estimate calculations and the datasets used in them.

| Method | Data sets used | Number of estimates |
| --- | --- | --- |
| Published estimates | Aebischer 2019 | 1 |
| Formal Release Records | APHAPR2019 data considering birds reported as 'held for release' | 1 |
| Estimates based on numbers bred and reared for release | a) APHAPR2019 data considering birds reported as 'held for breeding'<br>b) APHAPR2019 data considering birds reports as 'held for rearing'<br>c) DEFRA data on eggs/birds imported into the UK<br>d) Egg production values<br>e) Egg hatching proportions<br>f) Chick survival values | 2 |
| Estimates based on numbers of birds reported released on samples of shoots of differing sizes | a) GOPSOC2018 (Mean number of birds of all species per shoot = 4307)<br>b) PR data considering birds reported as 'held for releasing' (Mean number of birds per shoot = 3036 pheasants; 2887 red legged partridge; 754 mallard)<br>c) GOPDB2019 (Mean number of birds per shoot = 3238 pheasants; 1070 red legged partridge; 1816 mallard)<br>d) SSBS2017 (Mean number of birds per shoot with consideration of 3 shoot classes = 1532, 6212, 26241 birds/shoot)<br>e) Estimated numbers of shoots in the UK = 3300, 5000, 7300, 9100 (see Methods) | 16 |
| Estimates based on numbers of game keepers | a) NGO published estimate of numbers of keepers in the UK<br>b) SSBS2017 data on numbers of birds that each keeper can manage single handed and as part of a team | 2 |
| Estimates based on harvests from shoots | a) GOPSOC2018 (mean harvest/day = 98 birds)<br>b) GOPGS2018 (mean harvest/day = 112 birds)<br>c) GOPDB2019 (mean harvest/day = 165 birds)<br>d) SSBS2017 (mean harvest/day corrected for shoot class = 80, 148, 232 birds)<br>e) GOPSOC2018 (mean shoot days/season = 13)<br>f) SSBS2017 (mean shoot days/season corrected for shoot class = 9, 16, 41)<br>g) Estimated numbers of shoots in the UK = 3300, 5000, 7300, 9100 (see Methods)<br>h) Harvest efficiency (Robertson et al. 2017 = 33%; GOPGS2018 = 40%) | 32 |
| Estimates based on harvests by individual guns | PACEC2006 (number of people shooting released birds = 384,000)<br>PACEC2014 (number of people shooting driven game = 260,000)<br>PACEC2006 (number of gun days/season = total lowland shooting 2.32 million days; driven game only 1.5 million days)<br>GOPGS2018 (numbers of birds shot/gun/season corrected for spending group = 73, 185, 557, 111 birds/season)<br>Estimates of total gun days based on mean number of shoot days, shoot numbers and the size of gun teams, corrected for spending group<br>Harvest efficiency (Robertson et al. 2017 = 33%; GOPGS2018 = 40%) | 16 |
| Estimates based on BTO Breeding Bird Survey Data | BTO BBS data (Woodward 2020)<br>Mortality data (Pheasants = Madden et al. 2018; Red legged partridge = Hesford 2012) | 2 |

#### 1) Data Sets

##### 1) APHA Poultry Register (APHAPR2019)

Compulsory registration is required for individuals or organisations that breed, rear or release >50 gamebirds. Voluntary registration is available to those releasing <50 birds (Anon 2020a). During the registration process, registees are asked to report: Species (pheasant, partridge (no separation of red-legged and grey) and duck (no distinction by species); Livestock Unit Animal Production Usage (Shooting, Other); Livestock Unit Animal Purpose (breeding for shooting, rearing for shooting, release for shooting); and Usual Stock Numbers. I made a FOI request for this information on 29 Jan 2020 and received a response on 13 Feb 2020. There were 7902 records, but this does not correspond to 7902 separate locations because a single location may include all three species (three records) and/or up to three Animal Purposes per species. When filtered by Species and Animal Purpose (specifically release for shooting), we can be more certain that a single record relates to a single location and/or shoot and represents the number of birds of that species released there.

##### 2) Import Figures

Daniel Zeichner MP asked DEFRA how many pheasant and partridge (a) poults and (b) fertilised eggs were imported into the UK in 2019. They responded on 21 July 2020 (Prentice 2020) with details of import figures for both partridges (presumed red-legged partridge) and pheasants during 2019.

##### 3) Data extracted from Guns on Pegs advertising website (GOPDB2019)

Guns on Pegs (https://www.gunsonpegs.com/) is a commercial advertising site where shoots looking to let days or attract syndicate members can advertise. They can enter free-text descriptions of their shoot. Entry dates are not recorded but it has been operating for 7-8 years. Between January and March 2020, I read through descriptions of 697 lowland shoots in England that advertised shooting of pheasant, partridge (no attempt is made to distinguish red-legged from grey partridge) and ‘duck’ (not specified as mallard but often contrasted with ‘wildfowl’). I extracted data on the quarry species and the bag sizes offered at each shoot, and any information about the numbers of birds released, although only 22 shoots reported this. Shoots advertising on the Guns on Pegs website (and consequently also those that participate in the website’s surveys of shoot owners and gun clients (see below)) are likely to be a non-random sample of shoots and guns in the UK, with a bias towards larger commercial shoots.

##### 4) Guns on Pegs Game Shooting Census 2017 (GOPGSC2017)

Guns on Pegs conducts an annual survey of guns. The results from their 2017 survey are available https://2391de4ba78ae59a71f3-fe3f5161196526a8a7b5af72d4961ee5.ssl.cf3.rackcdn.com/1815/3011/2351/1110_0118_Guns_on_Pegs_Four_Page_Leaflet_Hayley_Clifton_Final_Web.pdf. The survey included responses from 12,143 guns and reported numbers of days shot per gun and daily bag size with separation of participants by their patterns of spend.

##### 5) Guns on Pegs Shoot Owner Census 2017 (GOPSOC2017)

This survey, also by Guns on Pegs, was conducted at the level of the shoot rather than individual guns and the results from their 2017 survey are available https://www.gunsonpegs.com/articles/shooting-talk/the-shooting-world-by-numbers-2018-season. The survey included responses from 652 shoots across the UK and reported numbers of days shot per season, numbers of birds released, numbers of birds shot providing measures of bag size and harvest efficiency and an estimate of the numbers of UK shoots.

##### 6) The Savills Shoot Benchmarking Survey (SSBS2017)

Savills and the Game and Wildlife Conservation Trust conducted a shoot benchmarking survey for the 2016/17 season and a summary of their findings is available https://www.gwct.org.uk/media/664264/shoot-benchmarking-example-participants-benchmarking-report-fictional.pdf. The survey included responses from 155 shoots and reported the numbers of shoots, numbers of birds released, number of days shot per season, and average bag sizes across three different size classes. Small shoots released up to 3,000 birds; Medium shoots released 3,000-10,000 birds and Large shoots released >10,000 birds.

##### 7) The Economic and Environmental Impact of Sporting Shooting/The Value of Shooting (PACEC2006/PACEC2014)

PACEC (Public and Corporate Economic Consultants) conducted surveys of shooting providers and participants in 2004 and again in 2011/12 (PACEC 2006, 2014), with responses from 2,096 in 2004 and 16,234 in 2011/12. These reported numbers of people engaged in game shooting and the total number of gun-days when game shooting occurred.

### 2) Estimating numbers of shoots in the UK

In order to make sense of the various mean values of numbers of days shot per season, mean bag size or mean numbers of birds released which could be obtained from the surveys described above, it is necessary to know how many shoots operate in the UK. This number is unknown, so I made a series of estimates to encompass a possible range of numbers that could then be used to calculate the numbers of birds being released.

The GOPSOC2017 reports, without any supporting information, that there are between 8000 and 10000 shoots in the UK. The PACEC2006 report states that 91% of shoots in the UK harvest released birds. Therefore, I corrected their reported figures to include only shoots where birds were released to gives estimates of between 7280 and 9100 sites.

A report by the Farm Animal Welfare Commission (FAWC 2008) provides an estimate of ∼7000 shoots in the UK that release pheasants (presumably including 300 that release partridges), derived from the opinions of various stakeholders with no further detail to substantiate this.

The APHAPR2019 indicates that 3307 sites hold pheasants for release, 1323 sites hold partridges for release and 578 sites hold duck for release. Given that few sites release only partridge or only duck, but that these generally all release pheasants, this suggests a minimum of 3300 sites release any birds for shooting.

Shoots of any appreciable size require a gamekeeper to manage them. Therefore, by knowing the number of gamekeepers in the country, it may be possible to estimate the number of shoots. The occupation of gamekeeper was last recorded in the 1981 census when ∼2500 were recorded (Tapper 1992). The National Gamekeepers Organisation (NGO) website reports, without any supporting information, that there are currently 3000 full time and a similar number of part-time gamekeepers in the UK (Anon 2020b). This number includes gamekeepers on grouse moors (perhaps 500 such moors (Anon 2016)) and wild-bird shoots (where releasing does not occur) and likely a small number of deer stalkers. However, I have used the original data in my further analyses. Any correction for gamekeepers not working on shoots that release birds will only reduce the estimated number of shoots in the UK. This ratio of full time to part time gamekeepers is supported by the Gamekeepers and Wildlife Survey that reported that of 965 gamekeepers participating, 52% were full time, 30% were amateur and 18% were part time (Ewald & Gibbs 2019) The PACEC2006 Survey reports the mean % of shooting providers employing various numbers of keepers. I excluded providers defined as ‘clubs’ which are described as shooting a variety of quarry on an area rather than the similar ‘syndicates’ which specifically shoot game. 33% of providers didn’t employ anyone. 24% employed one person part time. 24% employed one person full time. 19% employed more than one person full time (10% two or more; 9% three or more). These figures, suggesting that there are almost three times as many full time gamekeepers as part time ones and that about one third of people acting as gamekeepers are amateurs/unpaid, are similar to those reported by Ewald & Gibbs (2019) which gives us some confidence in them. These ratios from the gamekeeper survey and PACEC data, when combined with the absolute numbers reported by the NGO indicate that there are around 2000 shoots in the UK that have no paid gamekeeper (but each has an amateur gamekeeper managing their birds), a further thousand that employ a part time gamekeeper, a thousand that employ a full time gamekeeper and a further thousand that employ more than one game keeper. This suggests that there are around 5000 shoots in the UK that are managed by a gamekeeper. Some small shoots may be managed by individuals or groups of individuals who do not identify themselves as gamekeepers, even amateur ones. PACEC (2014) suggests that 91% of shoots release birds, so we might reduce the total accordingly, however, the ADAS (2005) report states that the NGO estimates that only 80-90% of gamekeepers are members. Therefore, I will not attempt to correct for either errors in membership or release behaviour and assume that the two errors cancel one another out so that the figure of 5000 shoots is reasonable.

Given the large uncertainty over the total number of shoots in the UK, I conducted four estimates for each shoot-level release measure These ranged from the 3300 shoots reported in the APHAPR2019, through the 5000 shoots that I estimate are active based on the numbers of gamekeepers in the UK to the reported (but unsubstantiated) 7000 (FAWC 2008) to 7300-9100 (GOPSOC2017).

### 3) Estimating the distribution of size classes of shoots in the UK

Shoots are not homogeneous in their structure or behaviours. Larger shoots may release more birds, shoot more days in the season and harvest larger bags on each shoot day. The SSBS2017 divides shoots into three classes based on the mean number of birds released, the average acreage shot, days shot per season and mean bag size. In their survey of 155 shoots, there are approximately equal numbers in each class, but given that larger shoots are more likely to be interested in their financial performance, I suspect that larger shoots are overrepresented in that survey. A more accurate reflection of the size class composition may be obtained by considering the distribution of shoots reporting release numbers in the APHAPR2019 following the definitions given in the SSBS2017. For pheasants, of the 3307 shoots reporting, 2,464 (75%) are classed as small; 543 (16%) are classed as medium; and 300 (9%) are large. For red-legged partridges, of the 1323 shoots reporting, 536 (41%) are classed as small; 344 (26%) are classed as medium; and 451 (34%) are large. For mallard, of the 578 shoots reporting, 413 (72%) are classed as small; 119 (21%) are classed as medium; and 45 (8%) are large. Therefore, considering all species, 63% shoots may be described as small, 21% as medium and 17% as large according to the definitions in the SSBS2017.

### 4) Estimating harvest efficiency

The harvest or bag size represents only a fraction of the birds that will have been released. Based on figures available for pheasants, it might be assumed that a third of released birds are harvested (Robertson et al. 2017). Other surveys have reported higher return rates, with the GOPSOC2017 reporting 40% return, and SSBS2017 reporting 38%. Consequently, in estimates that involved harvest figures, I multiplied them by three (assuming 33% harvest) or 2.5 (assuming 40% harvest) to estimate release sizes.

## RESULTS

I initially calculated 71 estimated release sizes to be compared against the published figure from Aebischer 2019 (Fig 2). The initial set of estimates ran from 10.5 to 177.7 million birds released annually across all species. These had a mean value of 41.2 million birds (95%CI 34.1-47.9 million) and a median value of 33.1million (IQR 21.4-49.6 million). However, these initial estimates were later refined (see below).

**Figure 2.**
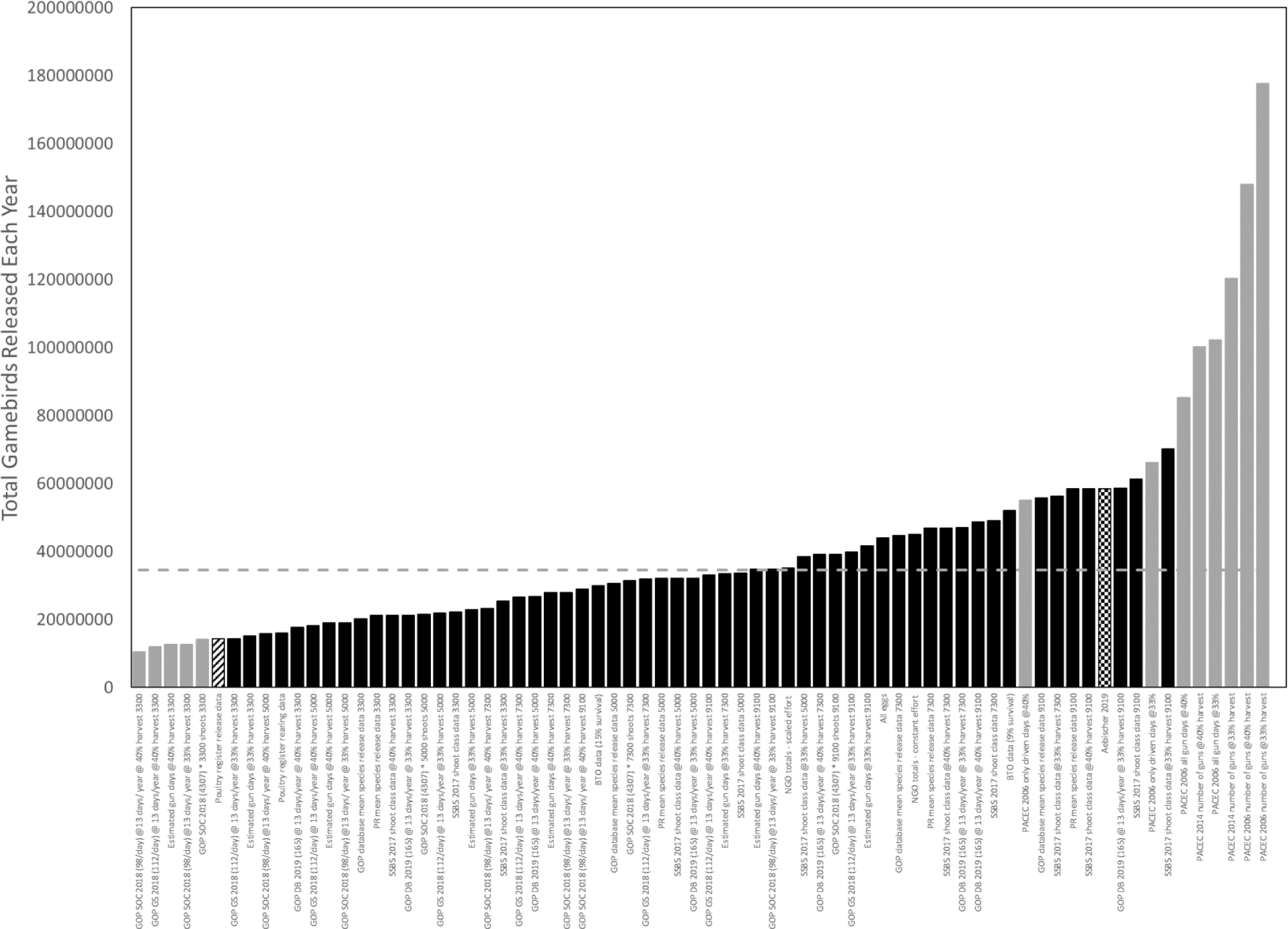
The 72 size-ordered estimates of the total numbers of gamebirds released in the UK. The diagonally hashed bar indicates the official figure derived from the APHA Poultry Register data for gamebirds held for release for shooting in the UK. The chequered bar indicates the figure published in Aebischer 2019. Grey bars indicate values that I consider to be uncredible, either because they fall below the official APHA Poultry Register figures or because they include data from the PACEC 2006 /2014 surveys that appears to be susceptible to over-reporting and bias. The mean value based on credible estimates is shown by the grey dashed line.

### 1) Published data estimating numbers of released birds (Aebischer 2019)

A mean estimate of 47 million pheasants and 10 million partridges released annually was calculated. Estimates of mallard release was not made. A total of 0.94 million mallard were reported shot and if we assume that half of these were released birds and that these represent one third of the mallard that were released (matching similar ratios for pheasant and partridges), then I estimate that 1.5 million mallard were released, giving a total of 58.5 million birds being released annually.

### 2) Poultry Register Data reporting numbers of birds held for release

The APHAPR2019 shows that: 10,039,379 pheasants and 3,819423 partridges and 435,907 duck (species unspecified) were reported as held for release for shooting. This produces a total of 14.3 million birds (of all species) released annually.

### 3) Estimates based on numbers bred and reared for release

Gamebird eggs and chicks are commonly produced outside the UK where more clement conditions stimulate higher and earlier productivity. Therefore, to estimate the numbers of birds that may be being bred for release it is necessary to consider numbers of eggs and chicks from UK and EU origins.

#### a) Birds bred in the UK based on breeding adults reported in the Poultry Register

There were 43,250 duck, 347,610 partridges and 1,052,436 pheasants registered as held for breeding in the APHAPR2019. Mallard and pheasants are both naturally polygynous, with a single male in the wild mating with several females. Game breeders take advantage of this situation to decrease the number of birds that they have to house (and feed) by placing several females with each male. This varies between breeders, but a ratio of one male to between seven and ten females is reported (Konteka et al. 2014), so we might presume that 88-91% of the birds registered for breeding are females. Using published productivity (57%) and hatchability (68%) figures (Datuin et al. 1996) for mallard I calculate that about 1.1 million mallard chicks could be bred for release. In the UK, pheasants used for egg production may be kept in cages or aviaries and these have implications for production rates and hatchability. Caged hens produce 29.9 chicks each while those in aviaries produce 9.8 chicks per hen (Konteka et al. 2014). It is unknown what proportion of the ∼947,000 hens are held in each housing condition, with Matheson et al. (2015) reporting that their use is currently rare but increasing and Canning (2005) stating that 5% of rearers were using such cages. Assuming that 25% of hens are housed in cages (with larger scale rearers investing in cages), then around 14.0 million pheasant chicks could be produced annually in the UK. Partridges vary in their productivity and egg hatchability with age (Mourão et al. 2010). Partridges are naturally socially monogamous and this mating system is replicated by breeders with partridges kept in pairs, so I estimate that about 6.8 million partridge chicks could be bred for release. In total, I estimate that ∼21.9 million gamebird chicks could be bred in the UK for release annually.

#### b) Birds bred outside the UK and imported as eggs or chicks

Records for 2019 show that 28,248,773 pheasant eggs were imported into the UK from Europe (Prentis 2020). Assuming hatching rates of 61% (Konteka et al. 2014), around 17.2 million chicks would hatch from imported eggs. 3,299,780 live pheasants and 1,673,165 live partridges were imported (Prentis 2020). From these figures, I estimate that up to ∼22.2 million birds that had been bred outside the UK could be released annually.

By combining the numbers of birds bred within and without the UK, I estimate that up to 44.1 million birds could be bred and released annually.

#### c) Poultry Register Data reporting numbers of birds reared

The APHA Poultry Register records do not consider a bird’s origin, and so records of birds reared for release could include both UK and international-bred birds. Records for 2019 show that: 11,196,463 pheasants, 4,607,688 partridges and 195,811 duck (species unspecified) were reported as reared for shooting. This produces a total of 16 million birds (of all species) that were reared and might be released annually.

### 4) Estimates based on numbers of birds reported released on samples of shoots of differing sizes

#### a) Using GOPSOC2017

This census reports a mean of 4,307 birds (with no error margins reported) being released on each of the 652 shoots that completed their survey. Assuming that the distribution of the surveyed shoots matches that in the general population (in reality it is likely that larger shoots are over represented in this survey - see above) then I estimate that between 14.2 and 39.2 million birds are released annually, depending on the number of shoots we assume to operate in the UK.

#### b) Using GOPDB2019

Release numbers from 22 shoots on the Guns on Pegs website indicated that, of the birds released on an average shoot, 73% were pheasants, 12% were partridges and 14% were mallard. For pheasants, the distribution of releases was very skewed with a mean of 3238 pheasants/shoot, but a median release size of 1500 birds/shoot with an IQ range of 1100-2500. For red-legged partridges, the distribution of releases was also very skewed with a mean of 1070 partridges/shoot, but a median release size of 400 birds/shoot with an IQ range of 275-1850. For mallard the distribution of releases was also very skewed with a mean of 1816 mallard/shoot, but a median release size of 300 birds/shoot with an IQ range of 150-575. Assuming that the distribution of the surveyed shoots matches that in the general population then I estimate (by multiplying mean releases per species and adding them together for each shoot) that between 20.2 and 55.7 million birds are released annually depending on the number of shoots we assume to operate in the UK.

#### c) Using APHAPR2019

For pheasants, the distribution of releases was very skewed with 6/3307 shoots reporting releasing >100,000 birds (111,000-200,000), a mean of 3036 pheasants/shoot, but a median release size of 850 birds/shoot with an IQ range of 400-2275. For red-legged partridges, the distribution of releases was also very skewed with 2/1323 shoots reporting releasing >100,000 birds (180,000 & 250,000), a mean of 2887 partridges/shoot, but a median release size of 500 birds/shoot with an IQ range of 200-2000. For mallard one record was for 150,000 birds released at a single shoot (of the 578 included). This was 25 times larger than the next largest value (6,000), so it could be a typo. If I retain this value in the calculations, there was a mean of 754 mallard/shoot, but a median release size of 200 with an IQ range of 100-500. Excluding the aberrant record reduces the mean to 495 mallard/shoot (but the median and IQ range are unchanged). Assuming that the distribution of the surveyed shoots matches that in the general population and using the lower figure for mallard, then I estimate (by multiplying mean releases per species and adding them together for each shoot) that between 21.2 and 58.4 million birds are released annually depending on the number of shoots we assume to operate in the UK.

#### d) Using SSBS2017

According to the SSBS2017, small shoots reported a mean release of 1,532 birds of all species; medium shoots reported a mean release of 6,212 birds; and large shoots reported a mean release of 26,241 birds. Assuming that of the shoots in the UK, 63% were small, 21% were medium and 17% were large (see above) then I estimate that between 22.2 and 61.2 million birds are released annually depending on the number of shoots assumed to operate in the UK.

### 5) Estimates of birds released on shoots given the available keepers to manage them

#### a) Assuming constant efforts across gamekeepers

The SSBS2017 reports that the mean number of birds released per full-time keeper is 10,204. Assuming that each full time gamekeeper can and does manage 10,000 released birds and that each part-time or amateur keeper can and does manage 5000 released birds, I calculate that with 3000 FT and 3000 PT gamekeepers [see section above], a total of 45 million birds could be released and managed.

#### b) Assuming scaling effects of gamekeeper efficiency

This assumption of constant efforts across keepers may be generous. The SSBS2017 reports that there are significant differences in the numbers of birds released per keeper depending on the shoot size, with full-time keepers on large shoots managing 11,478 each whereas those on medium shoots manage only 5,892 birds. Assuming that small shoots are all run by part-time and/or amateur keepers who are again half as efficient as a full time keeper on a medium shoot (thus managing 3000 birds each) and that the remaining full time keepers are equally split between medium and large shoots (see above), then I calculate that a total of 35.1 million birds could be released and managed.

### 6) Estimates based on harvests from shoots

Harvest levels depend on both the number of birds shot per day and the numbers of days shot per season. I obtained three estimates across all shoots for daily bag size. These ranged from 98/day (GOPSOC2017), through 112/day (GOPGS2017) to 165/day (GOPDB2019) in which 120 shoots offered for sale days with bags of < 100, 157 shoots offered days of 100-200 and 206 shoots offered days of >200. The GOPSOC2017 provided the only estimate of the mean number of days shot reporting 13 days/shoot which appeared to be stable over time with shoots anticipating the same number of shoot days in the following year. This approximates to shooting on one day per week during the three months of the main shooting season. I accounted for the harvest rate as described above to estimate release sizes. Therefore, I made a series of estimates of bird releases based on total harvest figures that included: harvest rate (33% or 40%) * mean bag size/day (98, 112 or 165) * number of days shot/season (13) * number of shoots in the UK (3300, 5000, 7300 or 9100). This approach produced estimates of between 10.5 and 58.6 million birds.

In addition to these estimates across all shoots, I obtained estimates for shoots of different sizes which differed in both the number of birds that they shoot per day and the number of days shot per season. Typically, larger commercial shoots both harvest larger bags per day and offer more days shooting per season. The SSBS2017 described small shoots as offering 9 days/season with a bag of 80 birds; medium shoots offering 16 days/season with a bag of 148 birds; and large shoots offering 41 days/season with a bag of 232 birds. Assuming that of the shoots operating in the UK (ranging between 3300 and 9100), 63% were small, 21% were medium and 17% were large (see above) and that harvest efficiency lies between 33% and 40% (as above), then I estimate that between 21.2 and 70.1 million birds are released annually.

### 7) Estimates based on harvests by individual guns

#### a) Based on the total number of guns in the UK

According to PACEC2006, 480,000 people shoot live quarry, with 80% of those shooting pheasant & partridges, suggesting that 384,000 people might shoot released birds annually. According to PACEC2014, 260,000 shoot driven game. The numbers of birds shot per gun varies and this variation is captured by the GOPGS2017. The survey identified three classes of gun from 6,510 individuals (of the 12,143 surveyed) who reported spend data. Like the shoots themselves, the number of birds shot per day typically increased with the number of days shot per year. Those described as ‘Low Spend’ comprised 2031 guns (17%) and shot a mean of 8 days/year with a mean bag size of 73 birds/shoot. Those described as ‘Medium Spend’ comprised 3759 guns (31%) and shot a mean of 12 days/year with a mean bag size of 123 birds/shoot. Those described as ‘High Spend’ comprised 730 guns (6%) and shot a mean of 21 days/year with a mean bag size of 212 birds/shoot. A further 5623 guns (46%) provided no spend data but reported that they shot a mean of 9 days/year with a mean bag size of 99 birds. I assume that a day’s bag was split equally between the typically eight guns in attendance, meaning that over one shooting season, a mean Low Spend gun shot 73 birds, a Medium Spend gun shot 184.5 birds, a High Spend gun shot 556.5 birds, and a No Spend gun shot 111.4 birds. I assumed that all those people reported as shooting birds in the PACEC survey fell into one of the four classes (Low, Medium, High, No spend data) and derived a series of estimates based on the total shooting population (384,000 or 260,000) * the proportions of each class (17%, 31%, 6%, 46%) * the mean season bag total of each class (73, 184.5, 556.5, 111.4) * harvest efficiency (see above: 33%, 40%). This approach produced estimates of between 100.3 and 177.7 million birds being released.

#### b) Based on the number of gun-days

An alternative approach to the total number of people shooting live quarry is to calculate the number of days that people shoot per season. Because the four different classes of guns described above shot different numbers of days/season as well as shooting different numbers of birds on each day, I calculated the percentage of shooting days that were taken by people of the four classes (Low = 13%, Medium = 35%, High = 12%, No spend data = 40%) and calculated the mean number of birds that the gun was likely to shoot on that day (Low = 9.13, Medium = 15.4, High = 26.5, No spend data = 12.4). The PACEC2006 reports 1.5 million gun days/season targeting driven lowland game and a further 820,000 days/season targeting walked up lowland game which may include released birds. This is a total of 2.32 million gun days/season with a conservative estimate of 1.5 million days if only shooting at driven and therefore presumed released birds is considered. Therefore, I calculated estimates of harvests based on these reported gun days (2.32 million, 1.5 million) being shared among the gun classes as described above and accounting for harvest efficiency (see above: 33%,40%). This approach produced estimates of between 55.1 and 102.3 million birds.

The reported figures from the PACEC survey seem high when considering the numbers of shoots likely to operate in the UK. Assuming that each shoot day involves 8 guns, then an average shoot would be expected to host between 21 days/season (with 9100 shoots in the country and a total of 1.5 million gun days) and 58 days/season (with 5000 shoots in the country and a total of 2.32 million gun days). These figures do not correspond to any other survey data and at the high end of the estimate require that every shoot be operating on almost 4 days a week throughout the main three months of the shooting season. Therefore, I also estimated more realistic numbers of gun days based on the mean numbers of days reported as being shot on each shoot (13 days/season) with 8 guns per day multiplied by the range of numbers of shoots in the UK (3300-9100). I then calculated estimates of harvests based on these estimated gun days (total gun days = 343200, 520000, 759000, 946400) being shared among the gun classes as described above and accounting for harvest efficiency (see above: 33%,40%). This approach produced estimates of between 12.6 and 41.7 million birds.

### 8) Estimates assuming that BTO Breeding Bird Survey Data is a remnant of the release population

An alternative to calculating release numbers from birds being shot, is to extrapolate backwards from the numbers of birds recorded as surviving following the shooting season and before the next year’s cohort has been released. 2,300,000 female pheasants were calculated to be present in GB during the 2016 breeding season (Woodward et al. 2020). Assuming that each hen has a single partner (generous, because pheasants are a polygynous breeding species with single males holding harems of several females, although other males may be classed as non-reproductive satellites), there are 4.6 million pheasants present in the breeding season. If these are the all the remnant survivors of the released birds and between 9 and 15% of released birds survive to the start of following breeding season (Madden et al. 2018), I estimate a release of 29-51 million pheasants. 72,500 red-legged partridge territories were calculated to be present in GB during 2016 breeding season (Woodward et al. 2020). Assuming that each territory comprises a single male and female, and that survival of released partridges to the end of January is 15% (Hesford 2012), I estimate a release of at least 0.96 million partridges. Wintering mallard in the UK include migrants so it is not possible to reliably attempt to calculate the size of mallard releases based on breeding populations. It was estimated that there were 665,000 mallard individuals present during winter 2012/13-2016/17 (Woodward et al. 2020), but these could include both wild and released birds. By combining values for pheasants and partridges (but excluding mallards due to unreliable data), I estimate that between 30 and 52 million birds could be released annually.

### Examination and Refinement of Estimate Data

Several of these estimates were clearly incompatible with one another. The minimum number of birds being released should be taken as the numbers reported in the APHAPR2019 (14.3 million) as it is highly unlikely that game managers would report releasing birds when they are not doing so. Five estimations fell below that value. Estimates that included data from the PACEC2006 and PACEC2014 that referred to the numbers of guns thought to partake in game shooting in the UK each year and the number of days that they shot each season led to some extremely high values. If 177 million birds were released annually then, if we assume that the average shoot releases 4307 birds, there would have to be 41,000 shoots operating in the country. Given that there are around 6,000 full and part-time keepers in the country, this seems unlikely. If we assume that there are 9100 shoots in the country (the maximum value used in my estimates) then each shoot would release 19,450 birds, which would provide 43 shooting days, each with a bag of 150 birds/day. The PACEC data regarding gun days also leads to high values. Assuming that each shoot day involves 8 guns, then, according to PACEC data, each shoot would be expected to host between 21 days/season (with 9100 shoots in the country and a total of 1.5 million gun days) and 58 days/season (with 5000 shoots in the country and a total of 2.32 million gun days). These figures do not correspond to any other survey data and at the high end of the estimate require that every shoot be operating on almost 4 days a week throughout the main three months of the shooting season. Consequently, I distrust the estimates based on PACEC2006 and PACEC2014 data and recommend excluding them.

When estimates falling below the APHAPR2019 values and those involving data from PACEC are excluded, there are 59 remaining estimates varying from 14.3 to 70.1 million birds released annually across all species. These had a mean value of 34.5 million birds (95%CI 30.9-38.1 million) and a median value of 32.1 million (IQR 22.0-44.9 million). These values have been estimated by combining across all species because data are often not provided per species. Therefore, to calculate the estimated numbers of each species being released, I took the proportions of each species reported as being released in the GOPDB2019 (73% pheasants, 12% partridges, 14% mallard) and the APHAPR2019 (70% pheasants, 27% partridges, 3% mallard) and multiplied them mean total bird values. This suggests that 24.3-25.3 million pheasants, 4.2-9.4 million partridges and 1.0-4.9 million mallard are released annually in the UK.

## DISCUSSION

The estimates of numbers of gamebirds being released for shooting in the UK that I derived were markedly lower than the most recent published estimate. The average value of my estimates of around 35 million birds of all three species is just 61% of the estimate of 57 million proposed by Aebischer (2019). That estimate falls outside either the IQR or 95%CI of my estimates, and my mean and median estimate falls outside his 95%CI. Just 5/57 (9%) of the credible estimates exceeded his estimate. They are also in stark contrast with estimates derived from the current formal means by which we might to obtain the measure, via the APHA Poultry Register. The 14.3 million birds recorded in the register are just 40% of my mean estimate and falls outside either the IQR or 95%CI of my estimates. They are just 25% of Aebischer’s (2019) estimate.

My estimates are highly variable. This is because the data on which they are based is sparse, often collected in an ad hoc fashion with little attempt to ensure that it is representative, and commonly relies on indirect modifiers such as measures of egg production or numbers of shoots that are themselves based on small and potentially biased samples. Some of the variation is due to the uncertainty over the numbers of shoots operating in the UK. With credible values ranging from 3,300 to 9,100 shoots, this introduces an almost three-fold variation in many estimates. Like bird releases, shoots are currently not required to register their existence. An accurate value is critical because many other values are derived at the level of the shoot including mean numbers of birds released and shot. Is it possible to increase our certainty about the number of shoots? I explained above how a consideration of the numbers and distributions of gamekeepers that work on the shoots might be used to refine estimates and that supports the value of around 5,000 shoots. An alternative approach is to consider the density of shoots in the UK and relate it to local observations on the ground. If we assume that gamebird release is concentrated in the lowlands and that of the 244,000km^2^ of the UK (World Bank 2020), 40% of UK is uplands (RSPB, no date), then there might be around 146,400km^2^ of land suitable to host released bird shoots. With 9,100 shoots, we might expect there to be one shoot every 16km^2^, with 5,000 shoots, one per 29km^2^ and with 3300 shoots, one per 44km^2^. Either use of remote sensing data or direct sampling on the ground in a stratified random sampling method is required to test which of these densities is most credible.

It is not simply the number of shoots that is likely to be critical, but also their size and activities. The distribution of releases across shoots appears to be very skewed. Both the APHAPR2019 data and the GOPDB2019 data reveal that there are many small shoots that operate over a relatively small area, shoot relatively few birds either on each shoot day or across the season as a whole because they only shoot on a few days, and consequently release few birds. There are also a small number of very large shoots that shoot large bags on many days, typically over much larger areas of land, and consequently release many birds. From the SSBS2017 classification of shoot sizes and based on APHAPR2019 release data, large shoots make up only 17% of UK shoots, yet they appear to shoot 63% of the birds. In the GOPSOC2017 it is reported of the distribution of their data that “just over 7% of shoots accounted for half of the total number of birds put down and shot.” In the APHAPR2019 data for pheasants, there were 6/3307 shoots reporting releasing >100,000 birds (111,000-200,000), but a median release size of 850 birds/shoot. For red-legged partridges, the distribution of releases was also very skewed with 2/1323 shoots reporting releasing >100,000 birds (180,000 & 250,000), but a median release size of 500 birds/shoot. Therefore, my estimates may not be especially sensitive to the total number of shoots in the country, but rather to the number of these especially large shoots. Given that such shoots are more likely to be commercial, they are also more likely to have advertised on Guns on Pegs website and hence have been included in the three datasets based on that website that I considered.

A second notable source of potential imprecision is the sparsity of human data covering the behaviour of those people rearing, releasing, managing and shooting the birds. The behaviour of guns and game managers is not homogenous and mirrors the skew seen in the scale of shoots described above, with the three groups of guns identified in the GOPGS2017 varying 7.5 fold in the numbers of birds that they shot each season, ranging from 70-550. Despite these values being derived from a large sample of >6000 people, the fact that it drew on people who had joined a website in order to purchase shooting suggests that they may be those individuals that shot more birds than most guns in a year. To refine these numbers, we require a deliberate sampling method that equally captures the behaviour of the occasional gun who is invited to walk around a friend’s farm twice a year, as well as that of the person for whom shooting is their main recreational hobby, shooting large numbers of birds on many days of the year at multiple sites, and can provide a reliable count of the numbers of such individuals.

The PACEC (2006 & 2014) reports did indeed attempt to collect such large and representative samples of gun and game manager behaviour, including survey results from >16,000 people. Yet the authors of the 2014 report state that even this sample was likely to underrepresent those with low levels of involvement or over represent larger providers of driven game shooting. Further, the PACEC surveys were framed as investigating economic, environmental and social benefits of shooting sports and as such, respondents may have been moved to over-report their activities in order to support an idea that their actions had these benefits. Such over-reporting, specifically in the numbers of people involved in shooting and of days that they shot each year may explain why my estimates based on those PACEC values consistently resulted in figures that seem incredible. Consequently, one explanation for the discrepancy between my own estimates and those of Aebischer (2019) is that he drew on data from the PACEC surveys reporting the numbers of birds (and other quarry) shot. If those data, like those regarding the shooting behaviour of respondents, were also imprecise and over-reported, then they may lead to especially high estimates of numbers of birds being released. However, even if I retain these estimates in my set, then my mean estimate of released birds (∼41 million) is still only 72% of that of Aebischer (2019).

A third indicator of likely imprecision in these measures are clear contradictions or anomalies when comparing data sources. For example, the data on egg imports (Prentis 2020) records ∼28 million pheasant eggs and no partridge eggs being imported, whereas Canning (2005) states that around 90% of red-legged partridges are hatched from imported eggs. One explanation is that some of the eggs described as being from pheasants are actually partridge eggs. A second example is that the APHAPR2019 records 15,999,962 birds being reared for released but 14,294,709 actually being held for release, a drop of 12% which is more than twice the reported mortality rate at this stage of <5% (Đorđevic *et al*. 2010). The fate of these missing birds is unknown. The APHAPR2019 also reports that 195,811 duck were held for rearing whereas more than twice as many (435,907) were held for releasing. It is unclear where these additional birds came from. These anomalies are all seen in official DEFRA records (of imports, poultry registration or background reports) which might usually be assumed to be the more reliable data sources. Consequently, although it may be desirable to base estimates of releases on the most credible data sources, it is unclear which these may be.

Because data reporting numbers of birds shot or their release and management did not commonly separate by species, it is difficult to be confident in the exact numbers of each species that are released. Pheasants are clearly the most commonly released bird, comprising >70% of the release population. They are also released a the most sites. In contrast, the numbers of partridge and mallard being released is less well known. This may be because some shoots specialise in these two species and so a small number of shoots that release large numbers of these species, indicated by the skewed APHAPR2019 and GOPDB2019 data, may mean that numbers of these species are especially sensitive to records from a small number of sites. Generally, the release patterns and broader ecology of partridge and mallard is less well understood compared to pheasants (Madden & Sage 2020) and a better understanding of these species is desirable in future work.

An attempt to verify these estimates is possible by setting the numbers of birds estimated to be released within the context of the rearers, shoots and broader landscape where they are bred, released, managed and shot. By considering how many birds we might expect to see being bred, released, managed or shot on an either an average shoot or a shoot that we can place in a particular class, future studies can judge whether an estimate is realistic. If large numbers of birds are believed to be released, then we should expect to also see large numbers of shoots operating over many days of a season with large harvests on each day. More detailed surveys of the numbers of shoots, their frequency and harvest intensity could be conducted either directly by observing at shoots, or via targeted surveys or even remotely by recording sounds of shooting, in order to establish how accurate these and future estimates may be. Given the uncertainty about the credibility of many of the components that contributed to my estimates, it is helpful to explore what these estimates may mean in terms of ‘the average shoot’. If my mean estimate of ∼35 million birds being released is correct, then assuming a 33% harvest rate, there are probably around 12 million birds being shot each year. If these are shared evenly across 9,100 shoots in the UK then an average shoot will harvest ∼1,300 birds per season. This could be achieved over 13 days of shooting per season with a daily bag of 100 birds. If there are fewer shoots, say 5,000 (based on the numbers of gamekeepers reported by the NGO), then an average shoot would harvest 2,400 birds per season, with either 24 days of shooting bags of 100 birds per day, equating to the shoot operating just under two days every week during the height of the shooting season, or about 10 days of shooting bags of 250 birds per day. If the estimate of Aebischer (2019) of 58.5 million birds (including mallard) being released is correct with ∼20 million being shot, then across 9100 shoots, an average shoot would harvest ∼2,200 birds per season, perhaps over 22 days shooting bags of 100 birds, or across 5000 shoots, an average shoot would harvest 4,000 birds. However, the data suggest that all shoots are not equal and instead there are a few large shoots and many smaller ones. Following the definitions of shoots considering their daily bag and numbers of days of shooting from SSBS2017, small shoots harvest 17.6% of the total bag, medium shoots harvest 19.4%, and large shoots harvest 63.0%. If my estimate is correct then a total annual harvest of 12 million birds, would suggest that there are 2,933 small shoots, 983 medium shoots and 794 large shoots, totalling 4710 shoots in the UK. If Aebischer’s estimate is correct then there should be 4,889 small shoots, 1,638 shoots and 1,325 large shoots, totalling 7,852 shoots in the UK.

Given such variability and uncertainty, is there any value in trying to make estimates? I would argue that even if my mean or range of estimates are inaccurate, they highlight that the existing measures derived from Aebischer (2019) or the official APHAPR2019 figures are also likely to be highly suspect because they do not appear to correspond to either the numbers of birds being bred, nor the activity patterns of game shoots, nor to the numbers of birds surveyed in the countryside during the breeding season. This has two implications. First, my estimate being higher than that of the APHA figures suggests that many game managers, perhaps most, are failing to comply with the register regulations. Second, my estimate being lower than that of Aebischer (2019) suggests that we are unlikely to have an accurate understanding of the ecological effects that the released birds and their associated management exert on the country’s wildlife and habitats (Madden & Sage 2020; Sage et al. 2020). Currently, the figure of 57 million gamebirds being released annually (excluding mallard) is one that is quoted by both advocates and opponents of gamebird release. For both ‘sides’ there is an incentive to report high figures: for opponents, such as the League Against Cruel Sports (LACS 2020), a large number of birds can be presented as both an increased source of direct negative environmental effects and a large pool of individuals that might be considered to suffer from poor husbandry or death; for advocates, high numbers can support high figures of employment (including that directed at habitat conservation) and economic benefits derived from game shooting. If my mean estimate is correct, being about 60% of the previously published figure, then it suggests that neither the scale of negative ethical or ecological effects of release, nor the positive economic benefits are as high as are currently assumed.

## Funding

This work was conducted during research time paid for by the University of Exeter.

## Conflicts of interest/Competing interests

JRM has no conflicts of interest to declare.

## Ethics approval

N/A

## Consent to participate

N/A

## Consent for publication

N/A

## Availability of data and material

All data are provided in the text or in accompanying references

## Code availability

N/A

## Notes

### Competing Interest Statement

The authors have declared no competing interest.

